# Transmorphic phage-guided systemic delivery of *TNFα* gene for the treatment of human paediatric medulloblastoma

**DOI:** 10.1101/2022.10.18.512650

**Authors:** Mariam Al-Bahrani, Paladd Asavarut, Sajee Waramit, Keittisak Suwan, Amin Hajitou

**Affiliations:** Phage Therapy Group, Department of Brain Sciences, Imperial College London, London, UK

**Keywords:** medulloblastoma, targeted therapy, systemic delivery, phage-guided delivery, tumor necrosis factor alpha (TNFα)

## Abstract

Medulloblastoma is the most common childhood brain tumor with an unfavorable prognosis and limited options of harmful treatments that are associated with devastating long-term side effects. Therefore, the development of safe, non-invasive and effective therapeutic approaches is required to save the quality of life of young medulloblastoma survivors. We postulated that therapeutic targeting is a solution. Thus, we used a recently designed tumor-targeted bacteriophage (phage)-derived particle, named transmorphic phage/AAV, TPA, to deliver a transgene expressing the tumor necrosis factor alpha (*TNFα*) for targeted systemic therapy of medulloblastoma. This vector was engineered to display the double cyclic RGD4C peptide to selectively target tumors after intravenous administration. Furthermore, the lack of native phage tropism to mammalian cells warrants safe and selective systemic delivery to the tumor microenvironment. *In vitro* RGD4C.TPA.*TNFα* treatment of human medulloblastoma cells generated efficient and selective *TNFα* expression, subsequently triggering cell death. Combination with the chemotherapeutic drug cisplatin, used clinically against medulloblastoma, resulted in augmented effect through the enhancement of *TNFα* gene expression. Systemic administration of RGD4C.TPA.*TNFα* to mice bearing subcutaneous medulloblastoma xenografts resulted in selective tumor homing of these particles, and consequently targeted tumor expression of TNFα, apoptosis, and destruction of the tumor vasculature. Thus, our RGD4C.TPA.*TNFα* particle provides selective and efficient systemic delivery of *TNFα* to medulloblastoma, yielding a potential TNFα anti-medulloblastoma therapy while sparing healthy tissues from the systemic toxicity of this cytokine.

## INTRODUCTION

Medulloblastoma is the most common malignant childhood brain tumor causing paediatric cancer-related deaths [1]. The current treatment regimen for medulloblastoma is surgical resection, radiotherapy and chemotherapy. Despite this aggressive treatment strategy, the survivors suffer from clinically significant disabilities and long-term medical complications such as endocrinological and neurocognitive deficits, as well as secondary tumors [2]. Resistance to chemotherapy is a major obstacle due, at least in part, to the blood-brain barrier (BBB); while high doses result in toxic side effects associated with lack of tumor selectivity [3]. Being a cerebellar tumor, the surgical resection on its own is associated with cognitive and motor deficits that are also induced by radiotherapy due to lesions in the cerebellum [4]. Therefore, development of novel therapeutic approaches that are non-invasive, tumor selective, safer, cost-effective and efficient is urgently needed to improve the current treatment strategy, avoid the long-term side-effects, and save the quality of life of these young survivors.

TNFα is an inflammatory cytokine that has been reported for its anticancer properties through induction of necrosis in certain tumor types, and apoptosis in others. Furthermore, TNFα was found to sensitize breast cancer cells to radiotherapy and chemotherapy through DNA damage and triggering necroptosis [5]. Despite these properties, TNFα has limited clinical application due to its high systemic toxicity and short half-life [6]. We postulated targeted gene delivery for selective production of TNFα in the tumor microenvironment as a solution to reduce systemic toxicity and control the time window of TNFα bioavailability. We previously reported targeted delivery of TNFα to tumors in experimental mouse models of cancer and in pet dogs with natural cancers [7, 8]. Recently, we designed a tumor-targeted transmorphic Phage/AAV (RGD4C.TPA) gene delivery system that we propose to apply for targeted *TNFα* therapy against paediatric medulloblastoma [9]. Cancer gene therapy offers a promising strategy by delivering therapeutic genes to the tumor microenvironment. As a tool for targeted and efficient gene delivery, bacteriophage-based vectors have proven their application in prostate and breast cancers, soft tissue sarcomas, gliomas, and pancreatic cancer [10-14] by using a hybrid phage vector, which was previously constructed by incorporating within the phage genome a mammalian transgene expression cassette flanked by inverted terminal repeats (ITRs) from the adeno-associated virus AAV2 [15]. Recently we reported a new generation of phage-based particles for targeted systemic gene delivery consisting of the packaging a recombinant rAAV2 DNA by coat proteins of a filamentous phage, named transmorphic phage/AAV, or TPA [9]. In the TPA particles, the phage genome was removed and an f1 origin of replication is the only remaining phage cis element to allow replication of the single stranded ITR-flanked transgene expression cassette in bacteria and its packaging into the phage capsid provided through infection by a helper phage [9]. The helper was modified to display the double cyclic RGD4C (CDCRGDCFC) ligand on its pIII minor coat proteins to specifically target tumors by binding to the α_v_β_3_ integrin receptor, and at a lower extent to α_v_β_5_ [16]. These integrin heterodimers are often overexpressed on tumor cells and tumor vasculature, but barely present in healthy tissues [16]. Beside targeted vector entry through integrin binding, transcriptional targeting is another strategy to target gene expression within cancer cells. This can be accomplished by controlling transgene expression using cancer-induced promoters such as the promoter of the glucose regulated protein 78 (*GRP78*), which is an inducible promoter that is activated by chemotherapy since the *GRP78* gene is associated with tumor resistance to treatment [16]. We previously reported that RGD4C/Phage-*GRP78*-guided gene expression is activated by the chemotherapeutic agent temozolomide (TMZ) in glioblastoma [16]. Other promoters such as the cytomegalovirus (*CMV*), is well characterized for its high transcriptional activity, which can be further enhanced by the chemotherapeutic drug cisplatin (CDDP), used clinically to treat paediatric medulloblastoma [17]. Therefore, combining chemotherapy with targeted gene delivery should achieve both an increase of expression of the therapeutic gene and reduce the dose of chemotherapeutic agents.

In this study, we used the recently reported TPA particles characterized by enhanced gene expression, high titer production and large DNA sequence accommodation[9], then constructed the RGD4C.TPA.*TNFα* (Fig. 1) to evaluate the efficacy of guided systemic delivery of *TNFα* gene for medulloblastoma treatment. We investigated the efficacy of RGD4C.TPA.*TNFα in vitro* using human medulloblastoma cell lines, and *in vivo* in tumor-bearing mice. We also explored targeted systemic *TNFα* therapy by RGD4C.TPA.*TNFα*, and its combination with the clinically used CDDP chemotherapeutic drug, as a strategy to enhance the therapeutic efficacy.

**Figure 1.**
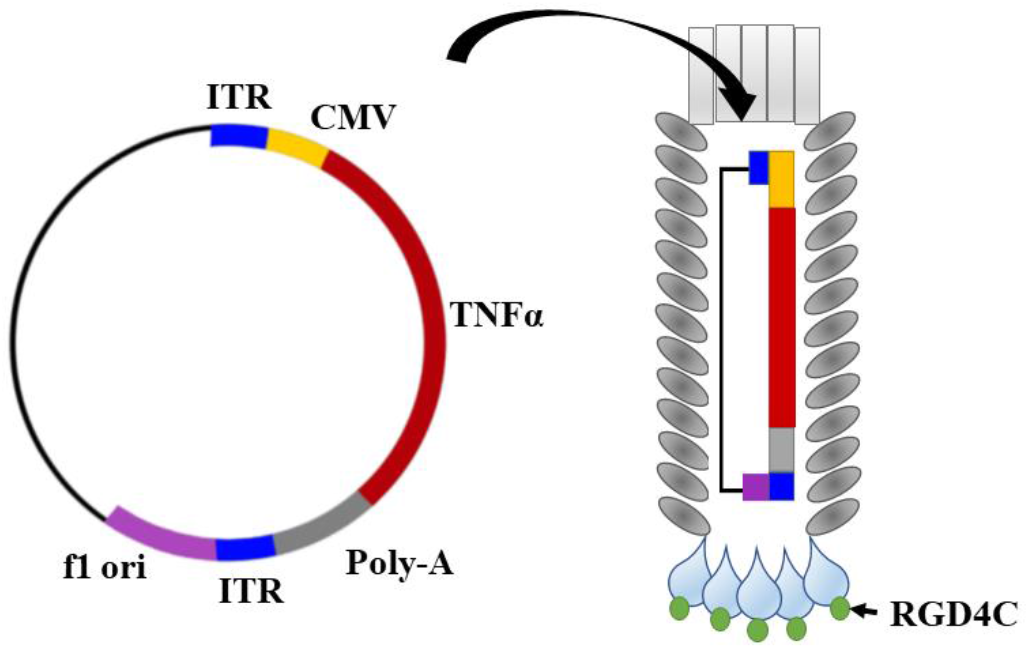
Schematic representation of targeted transmorphic phage/AAV (RGD4C.TPA) encoding *TNFα* gene. The RGD4C ligand is displayed on the pIII minor coat proteins of the TPA capsid. A *TNFα* therapeutic DNA sequence is inserted into a mammalian transgene cassette flanked by the ITR cis elements, from AAV2, and downstream a *CMV* promoter. A phage origin of replication, f1 *ori*, is the only phage genetic sequence present in the TPA DNA.

## RESULTS

### Human paediatric medulloblastoma cell lines express integrin cell surface receptors of RGD4C and are subsequently transduced by the RGD4C.TPA particle

The heterodimer α_v_β_3_ and α_v_β_5_ integrins have been linked to cancer cell growth and metastasis [18]. Binding of α_v_β_3_ and α_v_β_5_ to the extracellular matrix (ECM) proteins is known to be dependent on the presence of the RGD sequence on the ECM proteins [19]. Thus, UW228 and Daoy medulloblastoma cell lines were investigated for integrin expression and their suitability for targeted gene delivery by the RGD4C.TPA. The expression of α_v_, β_3_, and β_5_ subunits was confirmed by immunofluorescent staining to ensure that these cells express the α_v_β_3_ and α_v_β_5_ integrin receptors of RGD4C. As shown in Figure 2A, both UW228 and Daoy medulloblastoma cells express all the three integrin subunits.

**Figure 2.**
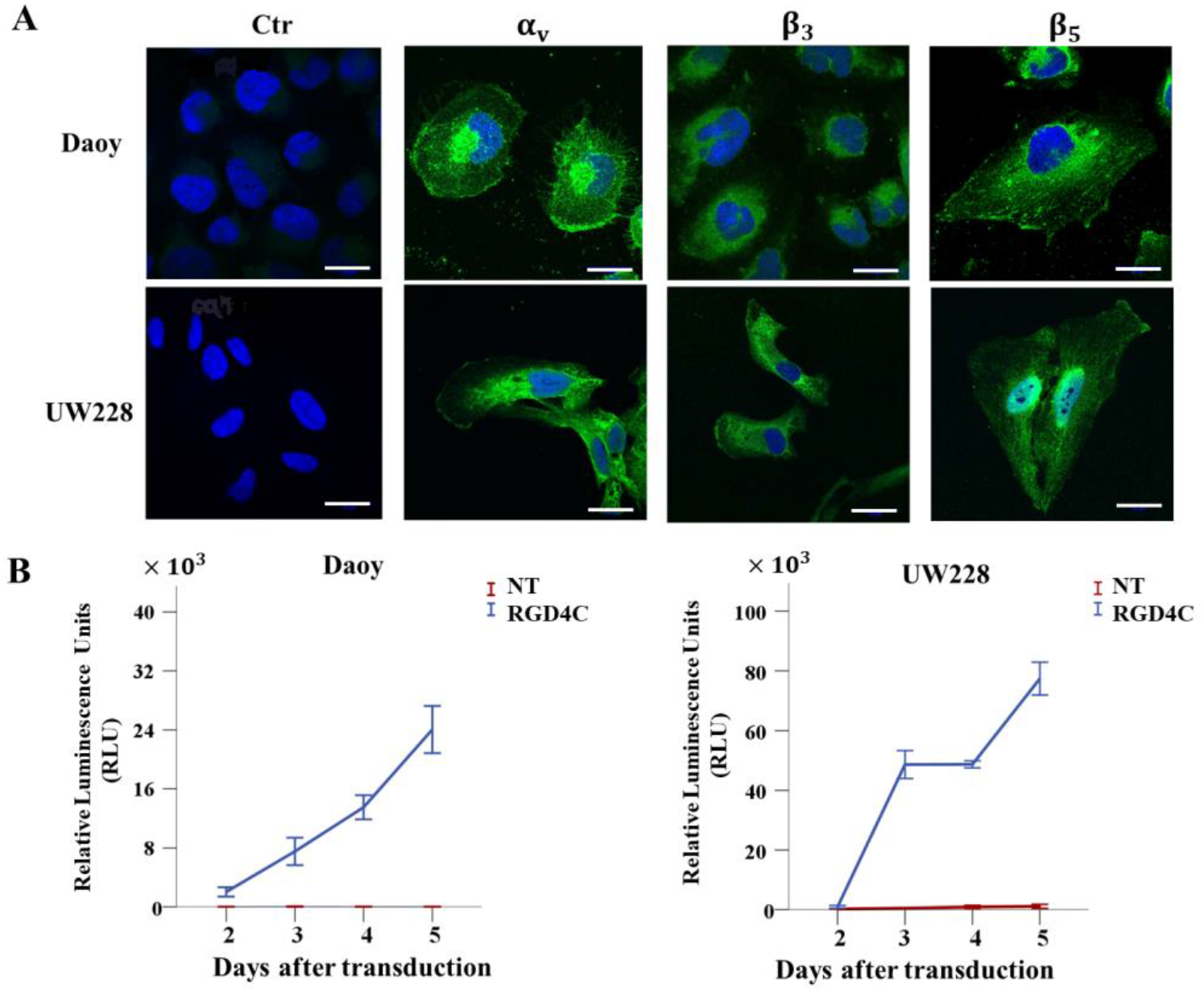
Expression of α_v_β_3_ and α_v_β_5_ integrin receptors in medulloblastoma mediates gene delivery by the RGD4C.TPA particle. **A**) immunofluorescent staining of medulloblastoma cells was carried without permeabilization using primary rabbit antibodies for α_v_, β_3_, and β_5_ integrins, followed by a secondary anti-rabbit IgG AlexaFluor-488 (Green) and 0.5μg/ml DAPI (4’,6-diamidino-2-phenylindole) (Blue). The control (Ctr) was stained with DAPI and secondary antibody alone. Scale bar, 25μm. **B**) Transduction of medulloblastoma cells with TPA.*Lucia* (targeted, RGD4C and non-targeted, NT). Secreted lucia was measured in the medium, daily and starting from day 2 after transduction then expressed as relative luminescence units (RLU).

Next, to evaluate gene delivery efficiency to medulloblastoma cells, we constructed and produced targeted RGD4C.TPA.*Lucia* carrying a reporter gene of the marine copepod secreted luciferase, *Lucia*. The use of secreted luciferase, lucia, is advantageous as it allows daily quantitative measurement of luciferase expression in the culture medium without lysing cells and provides higher sensitivity than the intracellular luciferase [20]. Tumor cells were transduced with targeted RGD4C.TPA.*Lucia* or non-targeted TPA.*Lucia* and expression of lucia was measured in the culture medium starting from day 2 post vector treatment (Fig. 2B). A marked increase in gene expression from the RGD4C.TPA.*Lucia* was observed overtime both in Daoy and UW228 cells; in contrast, no lucia expression was detected following cell treatment with the non-targeted TPA.*Lucia* particle. These data show that the RGD4C.TPA targets human medulloblastoma cells *in vitro* and generates selective gene expression dependent on the RGD4C ligand.

### RGD4C.TPA.*TNFα* targets TNFα production in medulloblastoma cells and induces tumor cell killing *in vitro*

After confirming the functionality of RGD4C.TPA delivery system using a reporter gene and its selectivity for the α_v_β_3_ and α_v_β_5_ integrins, we constructed vectors carrying a *TNFα* sequence as a therapeutic gene for medulloblastoma. To evaluate the RGD4C.TPA-mediated delivery of *TNFα*, we carried out analysis of gene expression both at the transcriptional level by real-time PCR and protein expression by ELISA to quantify the secreted TNFα in the media of transduced cells. Daoy and UW228 cancer cells transduced with RGD4C.TPA.*TNFα* showed high level of *TNFα* mRNA expression, which was undetectable in cells treated with the non-targeted TPA.*TNFα* (Fig. 3A). Then ELISA quantification of the secreted TNFα protein in the cell culture medium showed marked TNFα production from cells transduced with RGD4C.TPA.*TNFα*, but not cells treated with control non-targeted TPA.*TNFα* (Fig. 3B).

**Figure 3.**
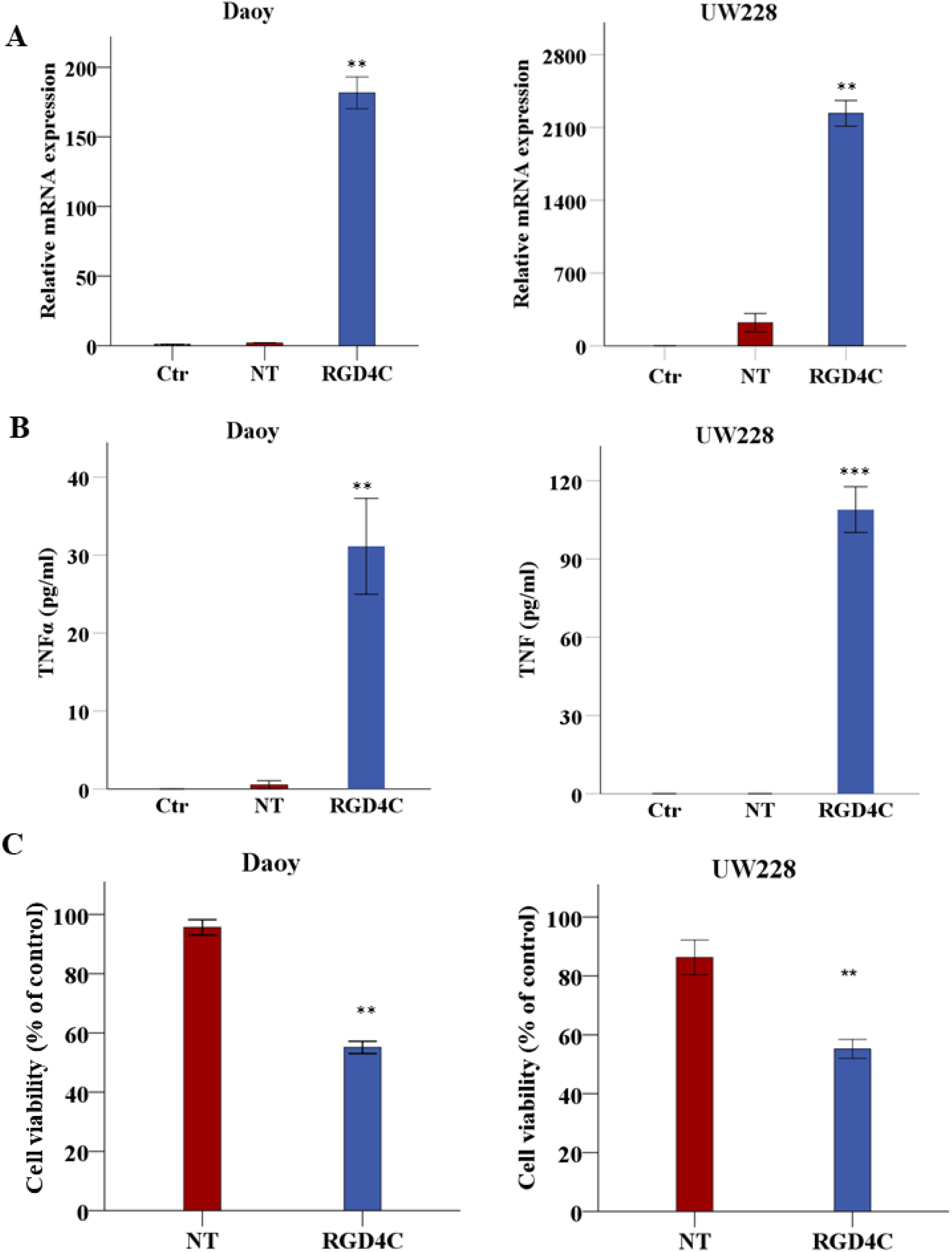
Targeted delivery of *TNFα* to medulloblastoma *in vitro* and assessment of the anti-tumor efficacy of RGD4C.TPA.*TNFα*. Daoy and UW228 cells were grown on multi well-plates, then transduced with RGD4C.TPA vector (RGD4C) carrying the *TNFα* transgene. Cells treated with the non-targeted vector (NT) were used as negative controls. **A**) Analysis of *TNFα* mRNA expression by qRT-PCR relative to *GAPDH* reference gene in Daoy and UW228 medulloblastoma cells at day 6 following treatment with targeted (RGD4C) or control non-targeted (NT) TPA. Control untreated cells (ctr) were also included in these experiments. **B**) TNF*α* protein production was measured by ELISA in the supernatant of medulloblastoma cells at day 6 post TPA transduction. **C**) Cell viability was measured at day 6 after transduction using sulphorhodamine B (SRB) assay and expressed as percentage of viability of untreated cells. Data are presented as mean ± SEM (standard error of the mean). **P≤0.01, ***P≤0.001. All experiments were repeated twice in triplicate and shown are representative experiments.

Next, we investigated the viability of medulloblastoma tumor cells following TPA transduction to assess the anti-tumor effect of RGD4C.TPA.*TNFα*. Cell death of Daoy and UW228 medulloblastoma was significantly induced by treatment with targeted RGD4C.TPA.*TNFα* as compared to the non-targeted TPA.*TNFα* which did not induce any significant tumor cell death (Fig. 3C). These findings show that medulloblastoma cell killing by RGD4C.TPA.*TNFα* is selective and mediated through the RGD4C tumor targeting ligand.

### Treatment with CDDP chemotherapy boosts TNF*α* expression from TPA and enhances medulloblastoma cell death

Our previous studies show that chemotherapy increases targeted gene therapy by phage-derived vectors against cancer including brain tumors [16, 21]. Therefore, we sought to test the effects of combining CDDP with RGD4C.TPA.*TNFα* particle, as this chemotherapeutic agent has been used clinically to treat medulloblastoma patients [22]. Thus, UW228 cells were transduced with RGD4C.TPA.*TNFα* under the control of the *GRP78* promoter (RGD4C.TPA.*GRP78*.*TNFα*) and selected with puromycin to generate stably transduced UW228.*GRP78*.*TNFα* tumor cells. Indeed, we previously reported that chemotherapy activates phage-mediated gene expression under the control of the *GRP78* promoter [16]. Cells stably transduced with a mock control TPA lacking the *TNFα* were also produced. Then, UW228.*GRP78*.*TNFα* were treated with 1μM and 5μM of CDDP and cell viability was measured over a time course at 16, 24, 48, and 72hrs post CDDP administration (Fig. 4A and B). CDDP treatment of UW228.*GRP78*.*TNFα* cells stably transduced with RGD4C.TPA.*GRP78*.*TNFα* induced higher cell killing compared to single treatment with CDDP or stable transduction with RGD4C.TPA.*GRP78*.*TNFα* alone.

**Figure 4.**
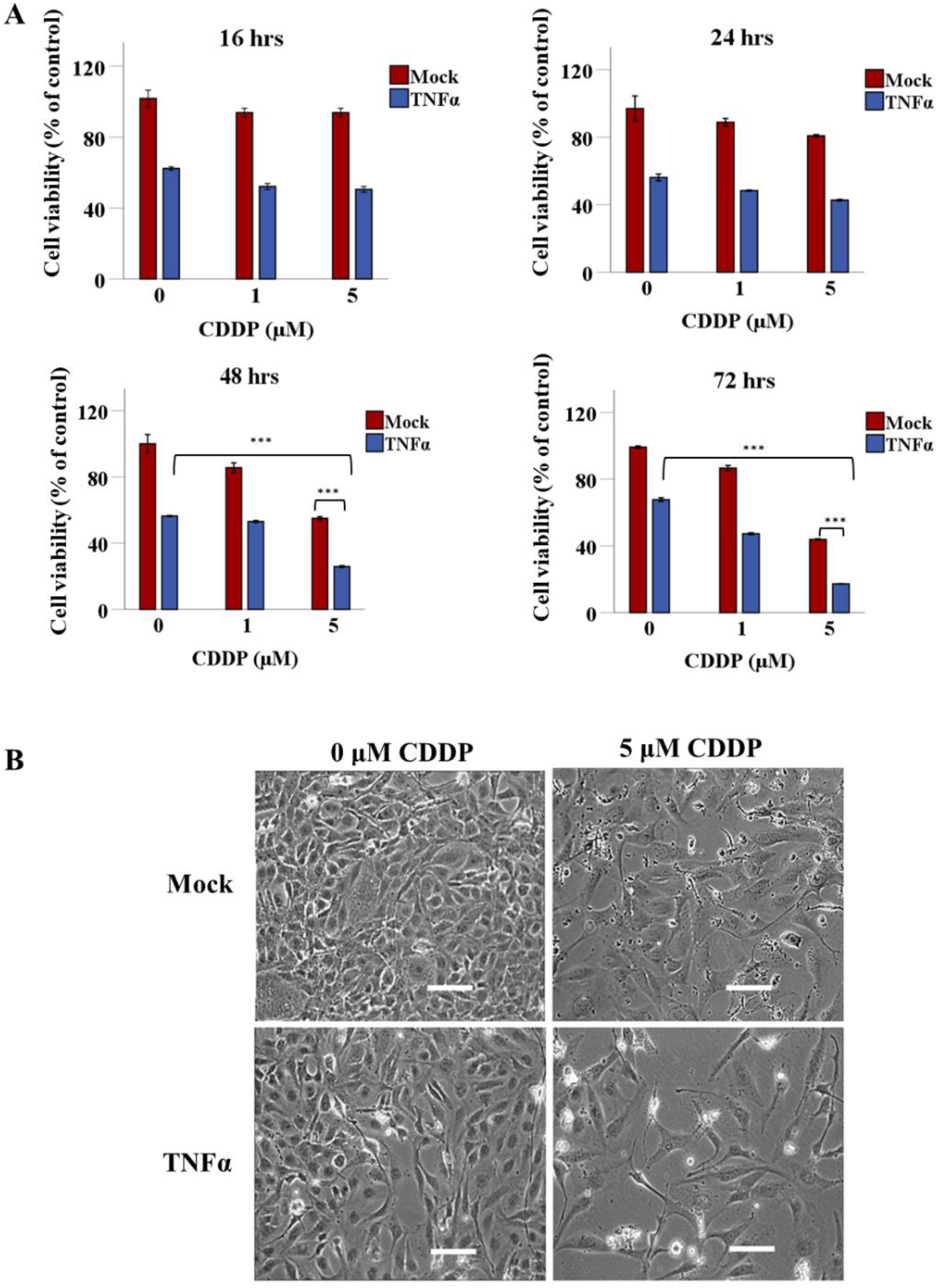
CDDP enhances TPA.*TNFα*-mediated medulloblastoma cell death. **A**) UW228 medulloblastoma cells carrying the *GRP78*.*TNFα* transgene expression cassette, were seeded in multi-well plates and allowed to grow for two days, then treated with 1 μM or 5μM of CDDP. Cells carrying the *GRP78* promoter but no *TNFα* gene (mock transduction) were also included as control. Cell viability was measured at 16, 24, 48 and 72hrs by using the SRB assay. **B**) Representative images of *GRP78*.*TNFα* tumor cells at 72hrs post treatment with 5μM of CDDP. Scale bar, 100μm. Data are presented as mean ± SEM (standard error of the mean). ***P≤0.001. All experiments were repeated twice in triplicate and shown are representative experiments.

Interestingly, CDDP treatment resulted in significant induction of TNFα expression under the control of *GRP78* promoter, where 1μM and 5μM CDDP-treated cells showed ∼2- and ∼7-folds increase in TNFα protein expression, respectively (Fig. 5A). This is consistent with our previous findings that TMZ chemotherapeutic drug, clinically used for brain tumors, activates the *GRP78* promoter in human glioblastoma [16]. Thus, before initiating *in vivo* studies, we sought to investigate the effect of CDDP on transgene expression by vectors carrying either *GRP78* or *CMV* promoters to select for the most suitable promoter to use in efficacy studies of TPA-guided *TNFα* expression in tumor-bearing mice. The *CMV* and *GRP78*-guided *Lucia* gene expression was tested using Daoy and UW228 cancer cells stably transduced with RGD4C.TPA.*CMV*.*Lucia* or RGD4C.TPA.*GRP78*.*Lucia*. Cells were then treated with two different concentrations of CDDP, 5μM and 10μM, and *Lucia* expression was measured at 48hrs post-chemotherapy treatment (Fig. 5B). Interestingly, lucia expression in 10μM CDDP-treated Daoy-*CMV-Lucia* cells significantly increased by ∼3.5-fold. While Daoy-*GRP78-Lucia* showed ∼2.4-fold increase with 10μM CDDP (Fig. 5B). These results were further confirmed in CDDP-treated UW228-*CMV-Lucia* and UW228-*GRP78-Lucia* cells, where 10μM CDDP showed ∼1.5-fold increase in *CMV*-guided lucia expression, and ∼1.8-fold increase in *GRP78*-guided *Lucia* expression at 48hrs (Fig. 5B). Therefore, based on the superiority of the *CMV* promoter in human medulloblastoma cells and its significant activation by CDDP, as compared to the *GRP78* promoter, we selected *CMV*-guided *TNFα* expression as the sequence to use in successive *in vivo* investigation of RGD4C.TPA.

**Figure 5.**
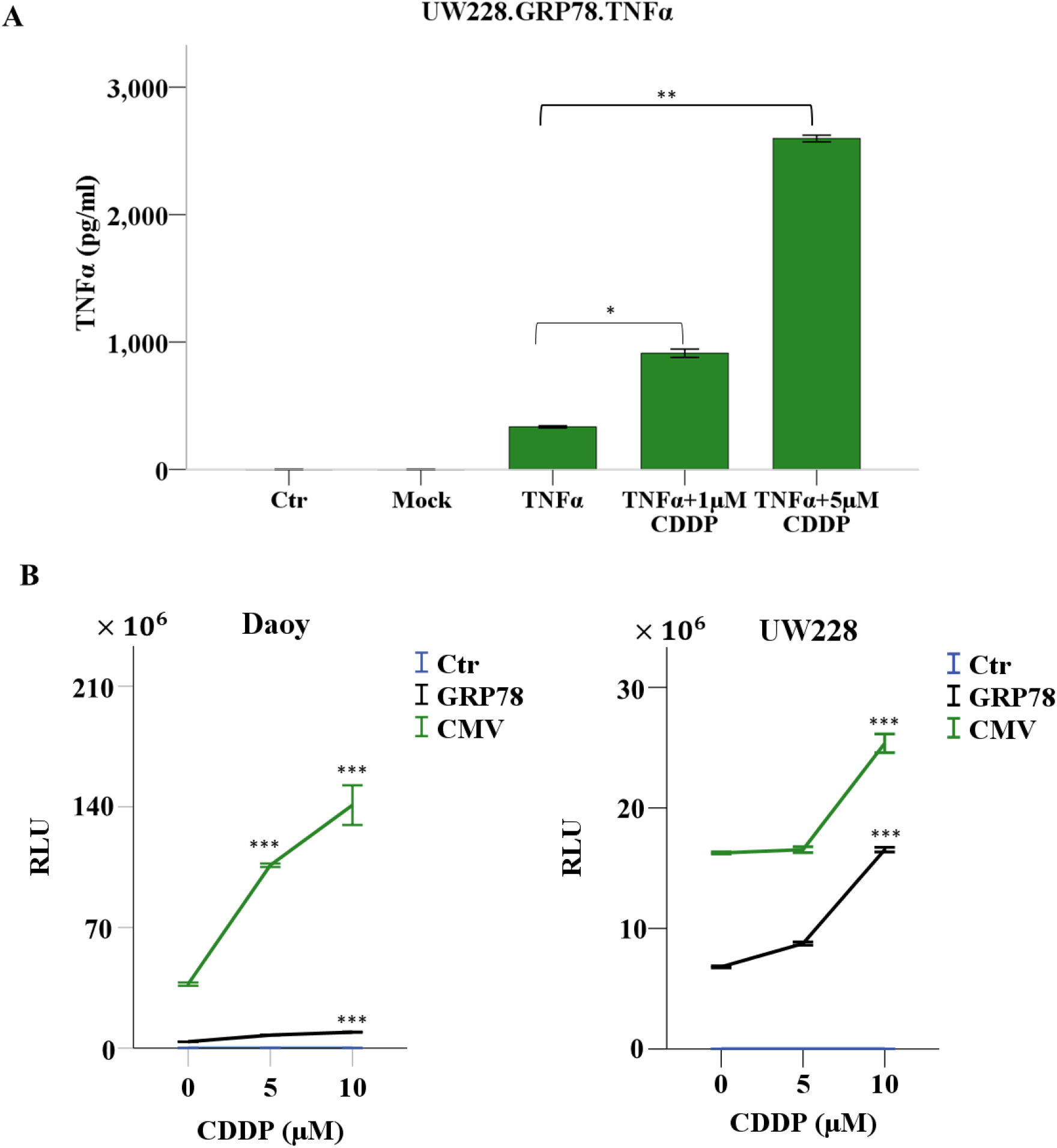
CDDP enhances gene expression under the control of *CMV* and *GRP78* promoters. **A**) Quantification of TNFα expression by ELISA following CDDP treatment for 72hrs of UW228 medulloblastoma cells transduced with TPA carrying *GRP78*.*TNFα*. Data were normalized to the percentage of viable control parental cells (ctr). Cells transduced with a mock vector lacking the *TNFα* transgene were also included as controls. **B**) Daoy and UW228 cells were treated with RGD4C.TPA.*GRP78*.*Lucia* or RGD4C.TPA.*CMV*.*Lucia* and selected with puromycin. The cells were seeded in multi-well plates and treated with CDDP. Secreted lucia expression was measured 48hrs after CDDP treatment and the results expressed as RLU. Data are presented as mean ± SEM. *P≤0.05, **P≤0.01, ***P≤0.001. All experiments were repeated twice. Shown are data from representative experiments.

### RGD4C.TPA.*TNFα* homing to medulloblastoma and biodistribution of *TNFα* expression in tumor-bearing mice following intravenous administration

Because efficacy and safety of *TNFα* gene systemic therapy will depend on its selective delivery to the tumor microenvironment, initial *in vivo* experiments were performed to investigate RGD4C.TPA-mediated *TNFα* gene delivery upon intravenous administration to tumor-bearing mice. As *in vivo* model, we established Daoy tumors stably expressing a firefly *Luciferase* (*Luc*) reporter gene to monitor tumor growth and response to therapy by using bioluminescent imaging (BLI) of the *Luc* reporter gene. Subcutaneous tumors were established in NOD/SCID mice by implanting *Luc*-labeled Daoy medulloblastoma cells. Next, upon detection of tumors in mice, the tumor-bearing mice were intravenously injected with 5 × 10^10^transducing units (TU)/mouse of targeted RGD4C.TPA.*TNFα* or non-targeted TPA.*TNFα* vectors expressing *TNFα* under the control of *CMV* promoter. First, we assessed the tumor homing of the RGD4C.TPA.*TNFα* particles upon systemic delivery. Thus, at 18hrs post vector administration, tumor-bearing mice were sacrificed, and the tumors were harvested. To assess vector localization within the tumors, immunofluorescent staining was carried for the phage capsid along with the blood vessel marker, CD31. The results revealed marked localization of the RGD4C.TPA.*TNFα* particles within the tumor tissue, and it was detected both in the tumor vasculature and tumor cells (Fig. 6A). No phage capsid staining was detected in tumors of mice following systemic treatment with the non-targeted TPA.*TNFα*.

**Figure 6.**
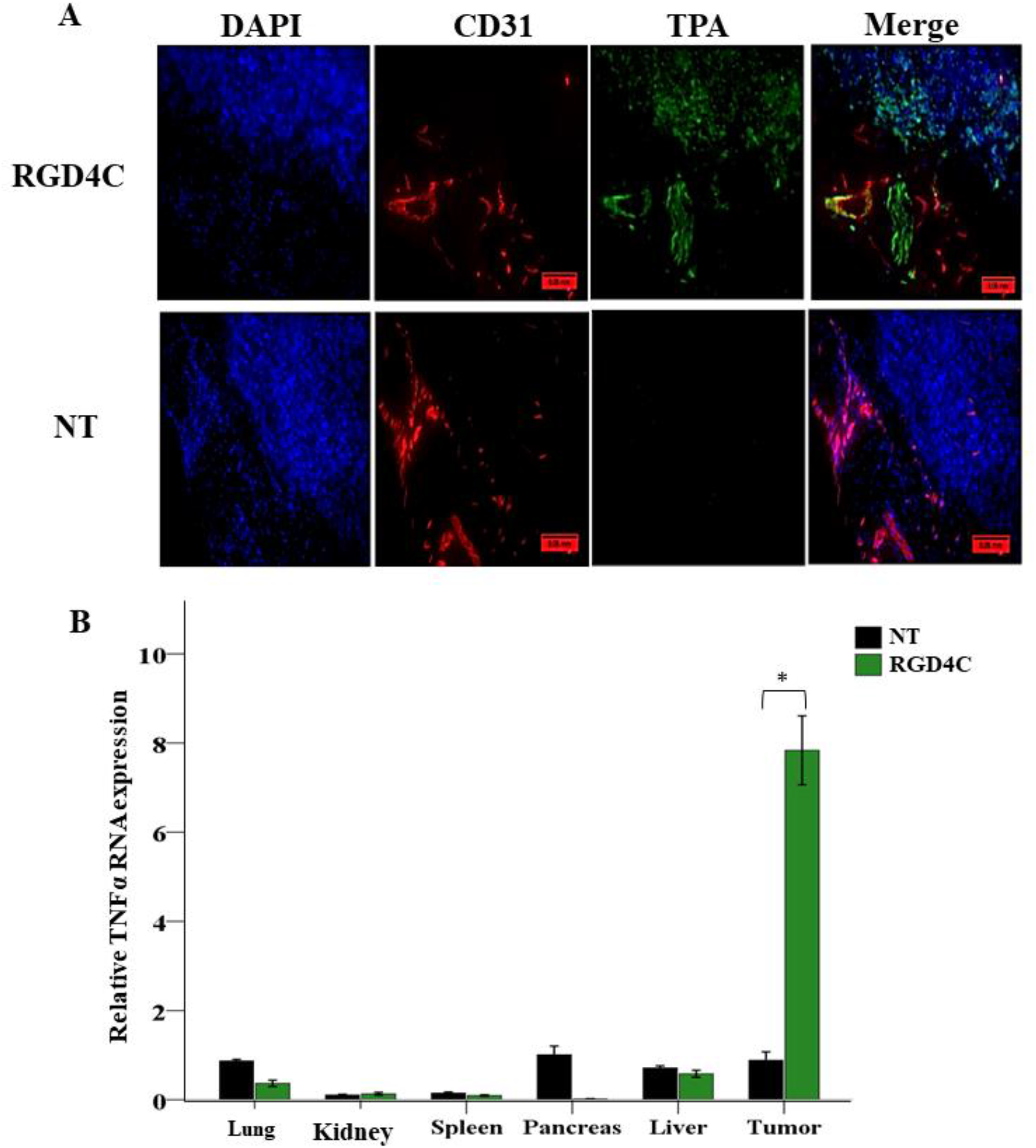
RGD4C.TPA.*TNFα* systemic targeting of medulloblastoma and biodistribution of *TNFα* expression. Daoy medulloblastoma cells were subcutaneously implanted into NOD/SCID mice and assigned randomly to either a control group TPA.*TNFα* (non-targeted) or targeted RGD4C.TPA.*TNFα* group. **A**) Tumor sections from treated mice were stained with primary antibodies for phage (green), CD31 (red) and 0.5μg/ml DAPI (blue). Scale bar, 50μm. **B**) RNA was extracted from tumors and various healthy tissues of tumor-bearing mice injected with 5×10^10^TU/mouse of targeted (RGD4C) or non-targeted TPA.*TNFα*. Expression of *TNFα* was determined by qRT-PCR relative to *GAPDH* reference gene. Data are expressed as mean ± SEM. *P≤0.05.

Next, to check whether the tumor homing of RGD4C.TPA.*TNFα* translates into selective expression of TNFα in medulloblastoma, tumor-bearing mice were intravenously injected with targeted or non-targeted TPA.*TNFα* twice at three days intervals. Mice were then culled at day 7 post vector delivery, and the tumors along with other major healthy tissues were harvested to perform qRT-PCR for analysis of *TNFα* gene expression (Fig. 6B). Interestingly, the data identified TNFα expression only in tumors harvested from RGD4C.TPA.*TNFα*-treated mice whereas no TNFα expression was detected in the lung, kidney, spleen, pancreas and liver (Fig. 6B). Additionally, no *TNFα* expression was detected in tumors or healthy tissues following intravenous administration of the non-targeted TPA.*TNFα* lacking the RGD4C ligand (Fig. 6B). Together these findings support the systemic tumor targeting of the vector and its efficacy for selective TNFα gene delivery to medulloblastoma.

### Targeted Systemic therapy with RGD4C.TPA.*TNFα* against medulloblastoma in mice

Next, we evaluated the therapeutic efficacy of systemic delivery of RGD4C.TPA.*TNFα* against medulloblastoma in mice. Once the subcutaneous tumors reached 200mm^3^ in size, non-targeted TPA.*TNFα* or targeted RGD4C.TPA.*TNFα* were injected intravenously twice, at days 0 and 4, via the vein tail (1 × 10^11^TU/mouse). A control group of mice injected with saline via the same route was also included. Moreover, additional groups of tumor-bearing mice received intraperitoneal CDDP (1mg/kg) administration along with the vector at days 0, 4, 7, 11, and 14 post vector treatment in order to investigate the effects of combination of chemotherapy and TNFα gene therapy, chemovirotherapy. This treatment regimen should allow expression of TNFα within the tumor, then administration of CDDP should induce activation of the promoter and further boost TNFα production. The tumor luminescence signals were monitored and evaluated overtime by bioluminescence imaging (Fig. 7A and B). Analysis of tumor growth showed that the tumors grew larger from day 4 to day 14, post TPA treatment, in groups of mice treated either with saline or control non-targeted vector (Fig. 7A); whereas mice receiving the targeted RGD4C.TPA.*TNFα* had their tumor growth reduced (Fig. 7A). These data were further confirmed by the loss of tumor luminescence signal, reflecting the tumor viability, following treatment with the targeted RGD4C.TPA.*TNFα* (Fig. 7B). Moreover, importantly, combination of RGD4C.TPA.*TNFα* with CDDP resulted in further antitumor effects, reduction in tumor size and tumor viability, when compared to RGD4C.TPA-*TNFα* or CDDP treated groups (Fig. 7A and B). We also monitored the animal weights during the course of therapy, and no weight loss was noticed (Fig. 7C).

**Figure 7.**
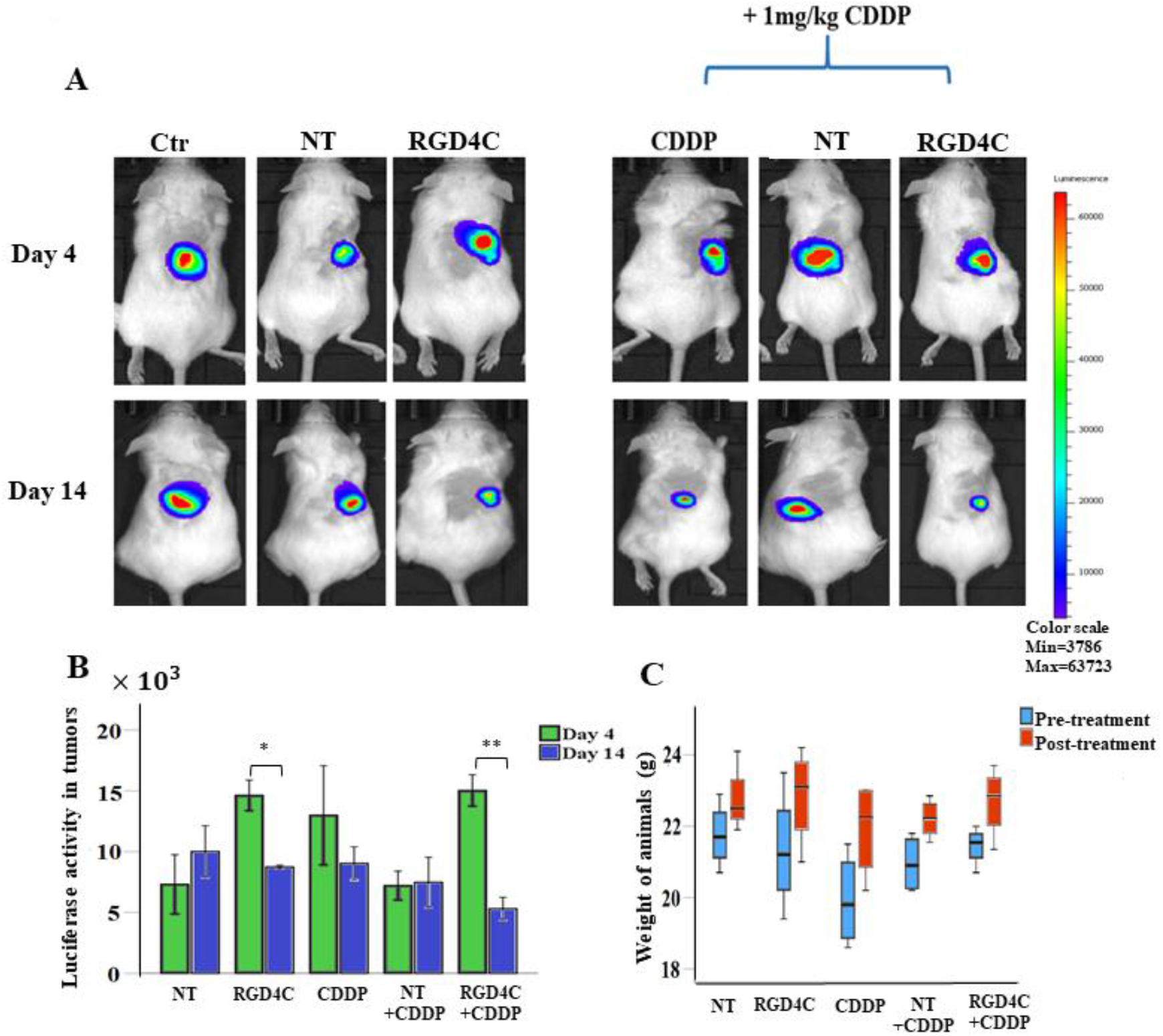
*In vivo* RGD4C.TPA.*TNFα* therapy in mice bearing subcutaneous medulloblastoma. **A**) Representative tumor-bearing mice showing *in vivo* bioluminescent imaging of *Luc* expression at days 4 and 14 post treatment. Control groups (Ctr) of tumor-bearing mice administered with saline or non-targeted (NT) TPA were also included. **B**) Tumor viability measured by bioluminescent imaging (P/sec/cm^2^/sr). **C**) Average animal weight during the experiment in each experimental group. The number of mice in each group is 4 (n=4). Data are presented as mean ± SEM. *P≤0.05, **P≤0.01

To further confirm the anti-tumor effects above, we carried out a comprehensive histopathological analysis of the tumors at the end of therapy. Using the terminal deoxynucleotide transferase dUTP nick end labelling (TUNEL) assay to detect apoptotic DNA fragmentation in tumor sections, we found that combination of RGD4C.TPA.*TNFα* with CDDP induced the highest level of apoptosis compared to CDDP or RGD4C.TPA.*TNFα* alone (Fig. 8A). Moreover, necrotic regions were observed in haematoxylin and eosin-stained tumor sections from mice treated with the combination treatment strategy (Fig. 8B).

**Figure 8.**
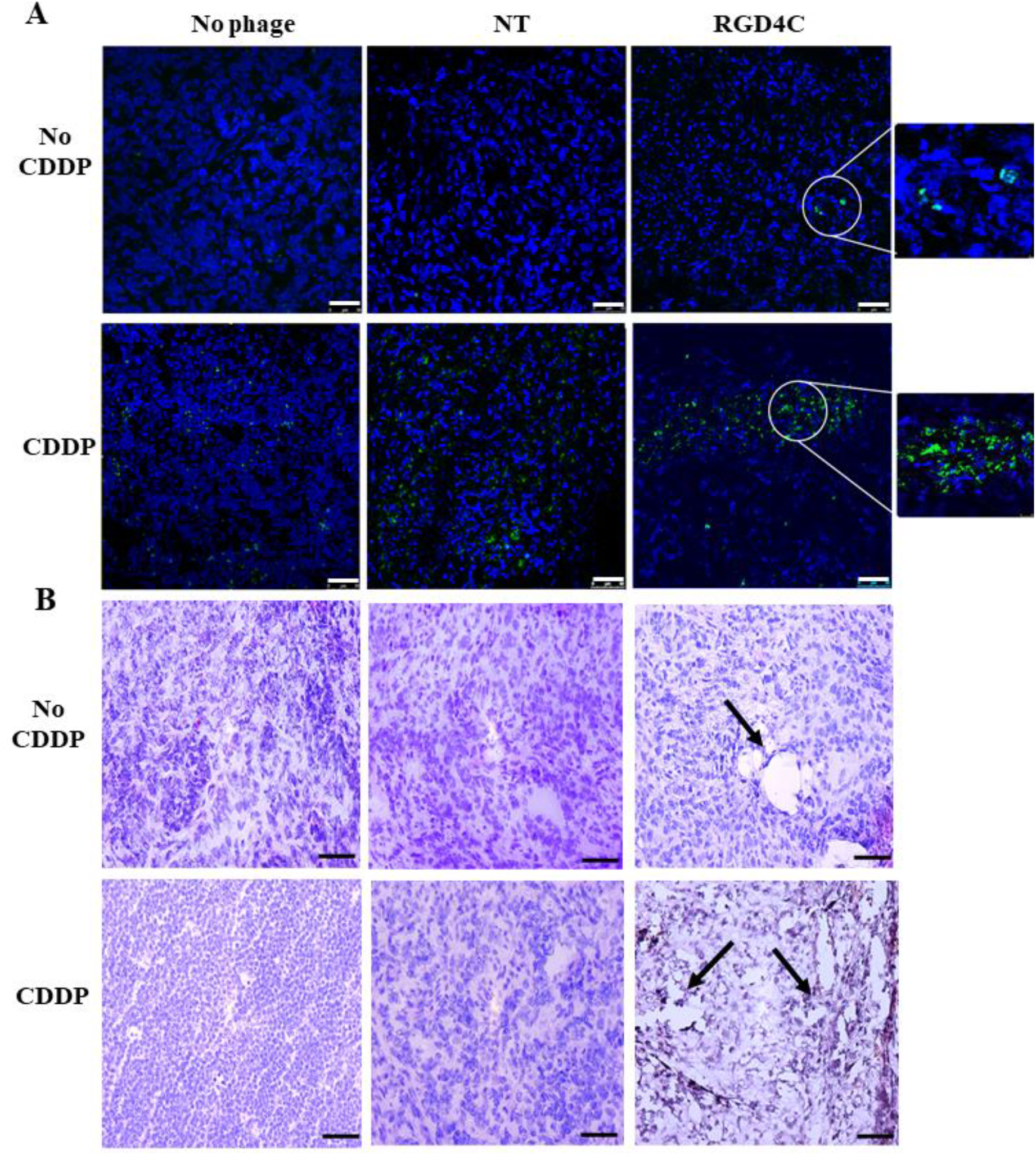
Histological analysis of tumors after therapy. **A**) TUNEL assay for detection of apoptosis in tumor sections. Post-treatment tumor sections were stained using the TUNEL assay and the images were taken using a confocal microscope. DAPI staining of the sections is shown in blue. Scale bar, 50μm. **B**) Tumor sections stained with haematoxylin and eosin. Arrows indicate necrotic regions in the tumor sections. The images were taken using a light microscope. Scale bar, 100μm.

Moreover, the tumor sections were stained with a CD31 antibody to investigate the therapeutic effect on the tumor vasculature since the RGD4C.TPA also targets the abnormal tumor blood vessels and TNFα was reported for its anti-angiogenic activity [6]. In contrast to the non-targeted TPA.*TNFα*, where the tumor vasculature remained intact (Fig. 9), the groups of mice treated with either RGD4C.TPA.*TNFα* or chemovirotherapy, showed extensive vascular damage as shown by the massive reduction in the tumor vasculature (Fig. 9).

**Figure 9.**
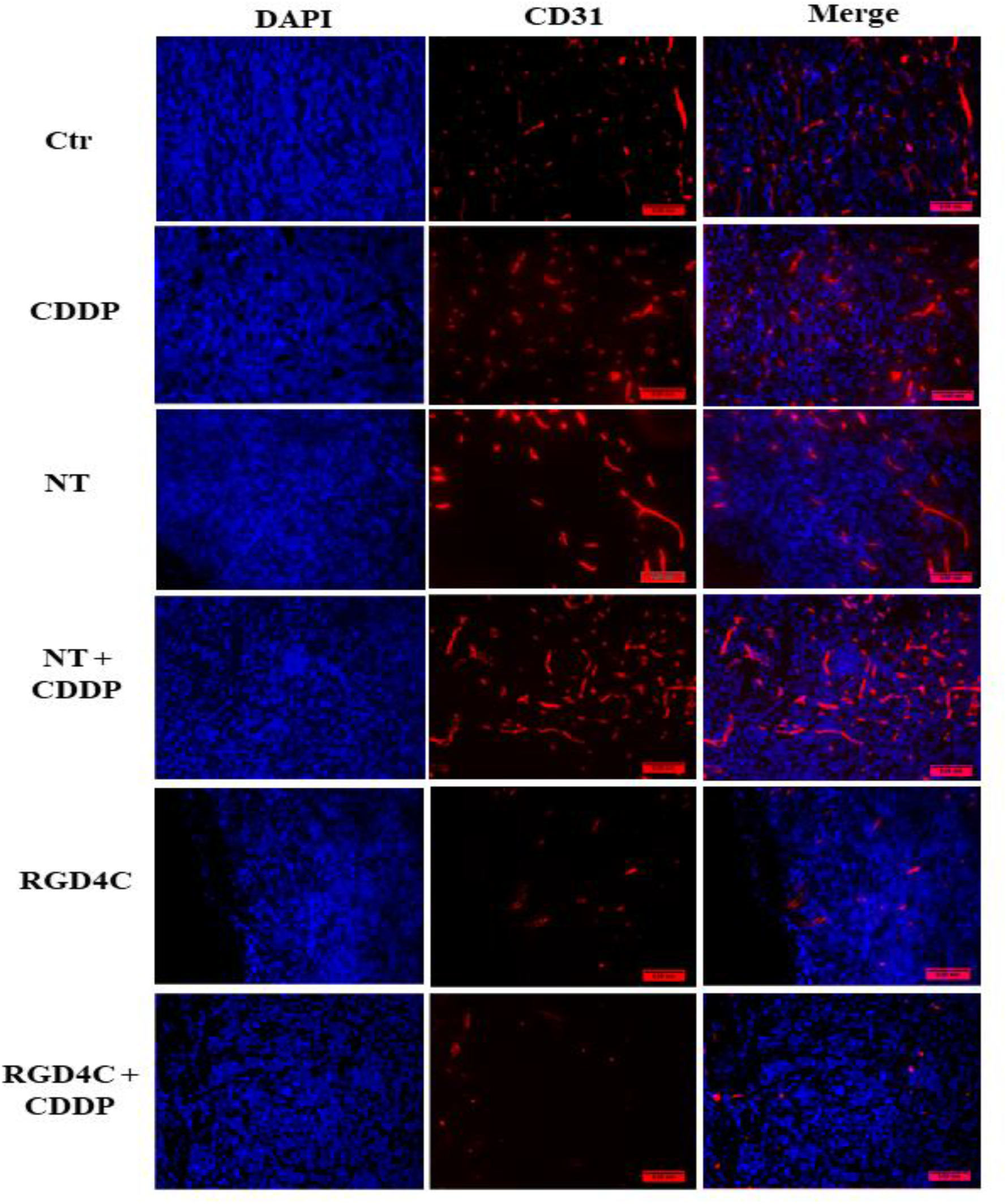
Immunohistochemistry analysis of the tumor vasculature. Tumor sections were stained with an anti-mouse CD31 (red), as a blood vessel marker, and DAPI for nucleus staining (blue). Scale bar, 50μm.

## DISCUSSION

In this study we investigated the efficacy of targeted systemic delivery of *TNFα* for the treatment of medulloblastoma by using a tumor selective RGD4C.TPA-mediated *TNFα* gene delivery platform. Phage-based vectors have proven to be advantageous for delivery to brain tumors because of their ability to cross the blood-brain barrier [16, 23-25]. Furthermore, the use of phage vectors is considered a safe therapeutic gene delivery strategy as they lack intrinsic tropism for mammalian cells and tissues, allowing their systemic administration [26]. The RGD4C peptide is one of the most characterized ligands for targeting the tumor endothelium as well as tumor cells [27]; hence our modified vector was genetically engineered to display the double cyclic RGD4C peptide on the phage capsid. Indeed, this peptide has been successfully used to target brain tumors such as glioblastoma [16, 28, 29]. Successful gene delivery to pre-clinical medulloblastoma models using mammalian viruses such as AAV has been reported to induce regression of tumor growth using Tis21 as a medulloblastoma suppressor gene; however, the direct delivery to the tumor tissue by intratumoral injections was inevitable [1]. Thus, to avoid repeated invasive local delivery strategies, systemic administration should be less invasive and safer when both tumor targeting and lack of mammalian tropism are combined into a single delivery system.

The use of *TNFα* in this study as a therapeutic gene showed efficient antitumor activity against medulloblastoma both *in vitro* and *in vivo* upon intravenous administration. Despite its anti-tumor effect, the clinical application of TNFα as a therapeutic agent has had limited success due to its severe systemic toxicity [6]. It is currently used in the clinic for the treatment of human melanoma and soft tissue sarcoma, but through local administration in the form of isolated limb perfusion [30]. Our phage gene delivery particle is designed to deliver the TNFα cytokine into the tumor microenvironment to ensure a local and dual selective effect both on the tumor microvasculature and tumor cells. Since this delivery model targets tumor cells and tumor vasculature, the expression of *TNFα* gene is localized and self-controlled; thus, once cell death is completed and the tumor is suppressed, *TNFα* gene expression should stop, consequently TNFα production ceases once the therapeutic effect is accomplished. Altogether the data show that our tumor-targeted phage delivery platform has the potential to bring systemic TNFα therapy back to the clinic to treat cancer patients via clinically non-invasive routes (e.g., intravenous).

Combining chemotherapeutic agents with gene therapy has long been used as a strategy to enhance therapeutic efficacy. Besides the additive effect of chemotherapy in inducing cell killing, it can induce gene expression through promoter activation. We previously reported such a strategy in which the chemotherapeutic agent temozolomide enhances gene expression in human glioblastoma through activation of the *GRP78* promoter [16]. Others have shown that CDDP enhances *CMV*-guided gene expression by inducing the CMV promoter activity [17]. Furthermore, we recently reported that the doxorubicin chemotherapeutic drug enhances phage-guided gene expression in glioma and melanoma cancer cells by facilitating vector trafficking into the nucleus of cells [21]. Besides the effect of chemotherapy and gene-guided expression, TNFα was shown to sensitize breast cancer cells to chemotherapy, including CDDP, and subsequently enhances the cytotoxic effect on cancer cells [5]. TNFα is also known to permeabilize the tumor vasculature when used in the form of isolated limb perfusion through down-regulation of VE-cadherin expression [31]. Consistently with this effect, our *in vivo* study showed that TPA-guided *TNFα* delivery resulted in tumor vascular destruction; such effect should enhance the efficacy of CDDP, allowing administration of reduced doses of chemotherapy.

Together our findings support our previous studies reporting chemotherapy as an adjuvant to activate RGD4C.Phage targeted cancer gene therapy [16, 21]. Additionally, our current findings have the potential to alter the clinical use of CDDP in medulloblastoma patients and should influence targeted systemic *TNFα* therapy in this type of brain tumors. Moreover, combinatorial treatment regimens represent a promising approach to overcome chemoresistance, which is a hallmark of medulloblastoma and should allow reduction of chemotherapeutic drug administration to less toxic and cost-effective doses.

## METHODS

### Construction of TPA expressing TNFα and Lucia transgenes

TPA constructs containing the RGD4C peptide were generated by inserting the *Lucia* (secreted luciferase) reporter or therapeutic (*TNFα*) genes into the TPA plasmid under the control of the cytomegalovirus, *CMV* promoter. Briefly, *Lucia* sequence flanked by HindIII and PmlI restriction sites was digested with HindIII and PmlI restriction enzymes. While, *TNFα* DNA sequence was flanked by EcoRI and BamHI restriction sites. The insert was then ligated with T4 DNA ligase (New England Biolabs, UK). The TPA particles were produced, as previously described [9], and purified from the culture supernatant of TG1 *E*.*coli* host bacteria, then filtered through 0.45μm filter. The TPA titer was calculated by infecting TG1 host bacteria with the TPA, the colonies were counted, and TPA titer was expressed as bacterial transducing units per μl (TU/μl).

### Cell culture

Daoy human medulloblastoma cell lines were purchased from the American Type Culture Collection (ATCC) and UW228 cell lines were provided by Dr. Jonathan Ham from the institute of Child Health, University College London. The cells were cultured in D-MEM medium containing 10% Fetal Bovine Serum (FBS), 2mM L-glutamine, 100U/ml Penicillin, and 100μg/ml streptomycin.

The cells were either authenticated by the supplier ATCC or by the collaborator who provided them. Upon receipt of these cell lines, they are first tested and cleared for mycoplasma contamination, then regularly tested for mycoplasma on a monthly basis by mycoplasma detection kit (Lonza, UK).

### Cell transduction by TPA

UW228 and Daoy were seeded in multi well plates to reach 70% confluency level after 2 days. Transduction medium was prepared with TPA particles, mixed with 30ng DEAE-Dextran per μg of TPA and incubated at room temperature for 15 minutes as we previously reported [32]. The growth medium was replaced with the transduction medium containing 1% FBS with either targeted RGD4C or non-targeted TPA and incubated for 24 hrs. The transduction medium was then removed and replaced with complete culture medium containing 10% FBS.

### Integrin immunostaining

Medulloblastoma cells were seeded on sterile glass coverslips and allowed to grow for two days prior to staining. The medium was discarded, cells were washed with phosphate-buffered saline (PBS) and fixed with 4% paraformaldehyde (PFA) at room temperature for 15 minutes, then washed and treated with 50mM ammonium chloride for five minutes. Next, cells were blocked with 2% bovine serum albumin (BSA) in PBS and 0.1% TWEEN20 at room temperature for one hour. Incubation with primary antibodies included anti-α_v_ and β_3_ and rabbit anti-β_5_ (Abcam, UK) diluted 1:50 in the blocking reagent overnight. Cells were then washed and incubated with the secondary Alexa Fluor 488-conjugated IgG antibodies (Invitrogen Life technologies), diluted 1:750 with 0.5μg/ml 4’,6-diamindino-2-phenylindole (DAPI) for 1 hour in the dark. The cells were washed and mounted with prolong gold antifade reagent (Life Technologies, UK). The samples were imaged using Zeiss LSM-780 inverted confocal laser scanning microscope.

### Luciferase reporter gene assay

Cells were seeded in multi-well plates and transduced with TPA-*Lucia*. The secreted lucia was measured in the culture medium using QUANTI-Luc reagent (InvivoGen, France), substrate for luciferase. Briefly, 10μl of the medium was mixed with 25μl of QUANTI-Luc. Luminescence was measured using GloMax Navigator (Promega, UK).

### Cell viability

Cells were washed with PBS and fixed with 10% trichloroacetic acid (prepared in serum free medium) overnight at 4°C. Fixed cells were washed with slow running tap water and allowed to dry at room temperature then stained with 0.4% sulphorodamine B (SRB, Sigma) for 30 minutes. Excess SRB dye was washed with 1% acetic acid, then cells were dried at room temperature. 150μl of 10mM Tris-Base solution (pH 10.5) were added and left on a shaking platform to solubilize the SRB. The absorbance was measured at 490nm using VersaMax Microplate Reader (Molecular Devices).

### Quantitative Real-Time PCR (qRT-PCR)

TNFα expression at transcriptional level was measured by real-time qRT-PCR using primer sequences specific to human *TNFα* and human *GAPDH* was used as reference gene (*TNFα:* forward, 5′CCCAGGGACCTCTCTCTAATCA; reverse, 5′AGCTGCCCCTCAGCTTGAG; *GAPDH*: forward, 5′CCCCTTCATTGACCTCAACTAC; reverse, 5′GATGACAAGCTTCCCGTTCTC). Total RNA was extracted with TRIZOL reagent (Ambion Life technologies, UK). 2μg of purified RNA was treated with DNaseI (Life technologies, UK), then heat inactivated after the addition of EDTA (final concentration 5mM) at 65°C for 10 minutes to stop DNaseI activity. cDNA was generated using GeneAmp RNA PCR core kit (Applied Biosystems, UK). SYBR green universal PCR master mix (Applied Biosystems, UK), and ABI7900 real-time PCR instrument (Applied Biosystems, UK). Gene expression was quantified using the comparative *C*_*t*_ method (Δ*C*_*t*_) and GAPDH as the reference gene. ΔΔ*C*_*t*_value was calculated against reference sample of non-transduced cells. The expression was calculated as 2^−ΔΔ*CT*^.

### Enzyme-linked immunosorbent assay (ELISA)

The level of TNF-α protein expression was evaluated after cell transduction with TPA-*TNFα* in the cultured medium using human TNFα ELISA kit (Biolegend, UK). First, the ELISA plates were coated with a capture antibody diluted in a coating buffer at 1:200 dilutions and incubated overnight at 4°C. Coating reagent was removed by washing 5 times with PBST (1X PBS with 0.05% (v/v) Tween-20) and blocked for one hour with 10% FBS. Then, the plates were washed, samples were added along with the standard and incubated for 2hrs with shaking. The wells were then washed, and detection antibody was added for one hour. This was followed by the addition of Avidin-HRP for 30 minutes. To minimize the background, the plates were washed and tetramethylbenzidine solution was added for 15 minutes in the dark. Finally, the reaction was stopped with 2N sulfuric acid and the absorbance was measured at 450nm using VersaMax Microplate Reader (Molecular Devices).

### Animal model and TPA-guided gene delivery

Human subcutaneous tumors were established in SCID immune-deficient female mice (Jackson laboratories) by inoculation of 15 × 10^6^ Daoy cells into the right flank of the mice. The tumors were allowed to grow to reach a volume of ∼200*mm*^3^. For the homing and biodistribution experiment, TPA particles (5 × 10^10^ TU/mouse) were administered twice intravenously into the tail vein three days apart. The mice were sacrificed at 18hrs after the second dose, then the tumors and organs were collected for further analysis [16].

For therapy experiments, 1 × 10^11^ TU/mouse TPA was injected intravenously at days 0, 4, 7, and 11. CDDP (1mg/kg) was administered through intraperitoneal route at days 7 and 11. Experiments involving living mice were carried out according to the Institutional and Home Office Guidelines, and under a granted Home Office-issued project license. The project license was first reviewed and approved by the Animal Welfare and Ethical Review Body (AWERB committee) at Imperial College London, before its final review and approval by the Home Office.

### Immunohistochemistry

TPA and tumor vascularization were immunostained on frozen sections using phage specific and CD31 primary antibodies, respectively. 6 μm tissue sections were cut from organs fixed in OCT. The sections were first fixed in ice-cold acetone for 10 minutes then washed with 0.1% PBST. Next, sections were blocked with 5% goat serum in PBST with 1% BSA for one hour at room temperature. The sections were incubated for 48hrs with rabbit anti-phage (1:500) (Sigma, UK) and rat anti-mouse CD31 (1:50) primary antibodies (BD Pharmingen) in PBST containing 2% goat serum. This was followed by washing with PBST and staining with the secondary antibodies, AlexaFluor 488 goat anti-rabbit IgG (1:500) and AlexaFluor 594 donkey anti-rat IgG (1:500) (Life Technologies, UK), as well as DAPI (1:3000) in 2% goat serum at room temperature for one hour in the dark. Finally, the samples were washed and mounted with ProLong gold antifade reagent (Life Technologies, UK).

For the evaluation of apoptosis in tumor sections, TUNEL assay was carried out using DeadEnd Fluorometric TUNEL System kit (Promega, UK). Briefly, the sections were fixed with 4% PFA, washed, and permeabilized with 20μg/ml proteinase K for 9 minutes. Then, the sections were washed with PBS, re-fixed with 4% PFA for 5 minutes and equilibrated with equilibrium buffer for 10 minutes at room temperature. Next, tissue sections were labeled with terminal deoxynucleotidyl transferase (TdT) reaction mixture for 1 hour at 37°C, then, immersed in saline-sodium citrate (SCC) for 15 minutes to stop the reaction. This was followed by washing with PBS and counterstaining with DAPI (1μg/ml) for 15 minutes. Finally, the tissue sections were washed and mounted with ProLong gold antifade reagent. Immunofluorescence images were acquired using Leica SP8 TCS confocal fluorescence microscope (Leica, UK).

### Haematoxylin and eosin staining

Tumor sections were fixed with 95% ethanol for 10 minutes, washed with tap water, and stained with haematoxylin for 5 minutes. Excess haematoxylin was washed for 5 minutes with tap water, followed by staining with eosin for another 5 minutes. The sections were washed again and dehydrated in a series of increased concentration of ethanol (70%, 90%, and 100%). Finally, the sections were cleared in xylene twice, mounted in DPX mounting medium and dried. The slides were examined under the light microscope (Olympus, Vanox, AHBT3).

### Statistical analysis

Statistical analyses were performed with IBM SPSS statistics 23. Error bars indicate the standard error of the mean (SEM). Significance was determined between groups using the student t-test and ANOVA for normally distributed data, and Mann-Whitney and Kruskal-Wallis as non-parametric tests. P values <0.05 was considered statistically significant and represented as follows: *P<0.05, **P≤0.01, ***P≤0.001.

## Acknowledgements

M.A. was funded by Kuwait University Ph.D. Program. This study was also supported by grants G0701159 and MR/T029226/1 of the UK Medical Research Council (MRC) to A.H., project grants from the Children with Cancer UK and Cancer Research UK to A.H and K.S. We also thank Grace Chu and Kaoutar Bentayebi at Imperial College London, for the technical assistance, and the Department of Brain Sciences for the equipment.

